# Transcriptome Profile Analysis of Twisted Leaf Disease Response in Susceptible Sugarcane with *Narenga porphyrocoma* Genetic Background

**DOI:** 10.1101/633081

**Authors:** Jinju Wei, Zhihui Xiu, Huiping Ou, Junhui Chen, Huayan Jiang, Xiaoqiu Zhang, Ronghua Zhang, Hui Zhou, Yiyun Gui, Haibi Li, Yangrui Li, Rongzhong Yang, Dongliang Huang, Hongwei Tan, Xihui Liu

## Abstract

Sugarcane is an important industrial crop with a high sugar yield that has become a leading energy crop worldwide. It is widely cultivated in tropical and subtropical regions. Various diseases beset the cultivation of sugarcane. The molecular study of disease resistance in sugarcane is limited by its complicated genome. In our study, RNA-seq was employed to detect the mechanism of twisted leaf disease tolerance in modern cultivar sugarcane, which derived from *Narenga porphyrocoma*. We completed high-throughput transcriptomic sequencing of 12 samples, including three stages of a susceptible (NSBC1_T “H3-8”) and an unsusceptible cultivar (NSBC1_CK “H-19”) with two biological repeats, respectively. Using the *Saccharum spontaneum* genome as reference, the average mapping ratio of the clean data was over 70%. Among the differentially expressed genes between H3-8 and H3-19, we focused on the analysis of hormone pathways and resistance (*R*) genes. The results showed that twisted leaf disease triggers hormone networks and around 40% of *R* genes conditioned lower expression in the susceptible cultivar. One of the possible reasons for H3-8 being susceptible to twisted leaf disease might be the null/retarded response of *R* genes, especially in pre-onset stage (46% down-regulated) of pathogens infection.

## Introduction

Sugarcane belongs to the Poaceae family, which includes maize, wheat, rice, sorghum, and many types of grass (Li 2010). Cultivated sugarcane is an important industrial crop with a high sugar yield and has become a major energy crop worldwide (Grivet and Arruda 2001; Yang et al. 2010). It is cultivated in ∼26 million hectares in tropical and subtropical regions of the world, producing up to 1.8 billion metric tons of crushable stems (Zhang et al. 2018). Sugarcane provides around 80% of the world’s sugar, with the secondary production as raw materials for pulp, ethanol and bioplastics (Lam et al. 2009; Li and Yang 2015). Modern sugarcane cultivars are interspecific hybrids which were generated from the cross between *Saccharum officinarum* and *Saccharum spontaneum*, followed by backcrossing into *Saccharum officinarum* to select sugar-poor relative traits (Roach 1972; Brandes and Sartoris 1936). In China, modern sugarcane is mainly distributed in the provinces of Guangxi, Yunnan, Guangdong, and Hainan. Some cultivars were also improved disease resistance by crossing with *Saccharum barberi Jesweit* or/and *Narenga porphyrocoma* (Gao et al. 2012; Liu, et al. 2012a; Liu, et al. 2012b).

Sugarcane diseases, such as smut, white leaf, and wilt/top rot/Pokkah Boeng,, are critical limitations of production, causing serious losses in yield and quality among susceptible cultivars (Hameed et al. 2015; Paulo et al. 2016; Su et al. 2016). Traditionally, the resistant traits of *Saccharum spontaneum, Saccharum barberi Jesweit* or/and *Narenga porphyrocoma* were introduced into *Saccharum officinarum* by hybridization to improve the disease resistance of cultivated sugarcane (Gao et al. 2012; Liu, et al. 2012a; Liu, et al. 2012b). With the development of breeding approaches, molecular breeding with precise genome information accelerates the collection of disease resistance genes in one cultivar. However, applying this to sugarcane is difficult due to its high polyploidy and complex genome, with ploidy levels ranging from 5× to 16×, and chromosome numbers from 2n = 40–128, with some even as high as 200 (Sreenivasan et al. 1987; Liu et al. 2012; Liu et al. 2012). The estimated polyploid genome size of sugarcane ranges from 3.36 to 12.64 Gb, and the monoploid genome size ranges from 760 to 985 Mb (D’hont et al. 1996; Zhang et al. which is larger than the rice (400 MB) and the sorghum (760 MB) genomes (Soderlund et al., 2011). Such complex genetic background blocks the sequencing of the whole sugarcane genome. Studies on biological traits, such as biomass yield, sugar accumulation, and stress tolerance, have focused on transcriptome analysis (Kido et al. 2012; Fracasso et al. 2016; Huang et al. 2016). The transcriptome reveals specific transcripts produced under biotic stress (e.g., smut) and abiotic stress (e.g., drought) conditions in sugarcane (Iskandar et al. 2011; Mattielllo et al. 2015).

With the *Saccharum spontaneum* genome, which is relative high quality, as reference (Zhang et al. 2018), we employed transcriptome profiling to analyze the genes and pathways involved in twist leaf disease in modern sugarcane cultivar. Twisted leaf disease, caused by *Phoma* sp., which is one of the largest fungal genera, was first reported in Guangxi, China, in 2014, when more than 5% of sugarcane was infected in the field. Twisted leaf disease is somewhat similar to Pokkah Boeng disease (caused by *Fusarium moniliforme* Sheldon). The symptoms begin with yellowing on the midribs and leaf margins, then spread to the entire leaf, along with twisting and curling of the crown leaves (Lin et al. 2014). The modern sugarcane cultivar used in this study was generated from the BC_1_ generation offspring of *Narenga porphyrocom*a via crossing and backcrossing with cultivated sugarcane varieties, so that it harbored the *Narenga porphyrocom* genetic background. The RNA-seq (two biological repeats) comparison between un-susceptible and susceptible cultivars in different stages of infection provided a reference for understanding the mechanism of twisted leaf disease. Analysis revealed that twisted leaf disease triggered the hormone network. *R* genes expression profile showed ∼40% *R* genes expressed lower in susceptible cultivar. The null/retarded response of *R* genes might be one of the possible reasons for H3-8 being susceptible to twisted leaf disease.

## Results

### RNA Sequencing and mapping to the reference genome

Compared to the un-susceptible cultivar, the susceptible cultivar displayed twisted leaves after infection with the disease (Fig. 1a). To obtain the gene expression profile associated with twisted leaf disease in susceptible sugarcane, twelve sequencing cDNA libraries were constructed. They contained a susceptible genotype (NSBC_1__T) and un-susceptible genotype (NSBC_1__CK) at three stages (pre-onset, early, and serious symptoms) (Fig. 1b) with two biological repeats (2 cultivars ×3 stages ×2 biological repeats). As shown in Table 1, a total of 258,301,460 and 279,952,436 raw reads were generated from the NSBC_1__T and NSBC_1__CK library by sequencing on the Illumina Hiseq 2000 platform, respectively. After filtering low-quality reads, unknown nucleotides, and the contained adapters, 251,482,172 and 273,757,964 clean reads were left in the NSBC_1__T and NSBC_1__CK libraries, respectively. Using the genome of *Saccharum spontaneum* as reference (Zhang et al. 2018), we mapped the clean reads to the reference via HISAT (hierarchical indexing for spliced alignment of transcripts) (Kim et al. 2015). All samples conditioned over 70% mapping ratio (average 76.74%). The details were summarized in Table 2.

**Table 1.**
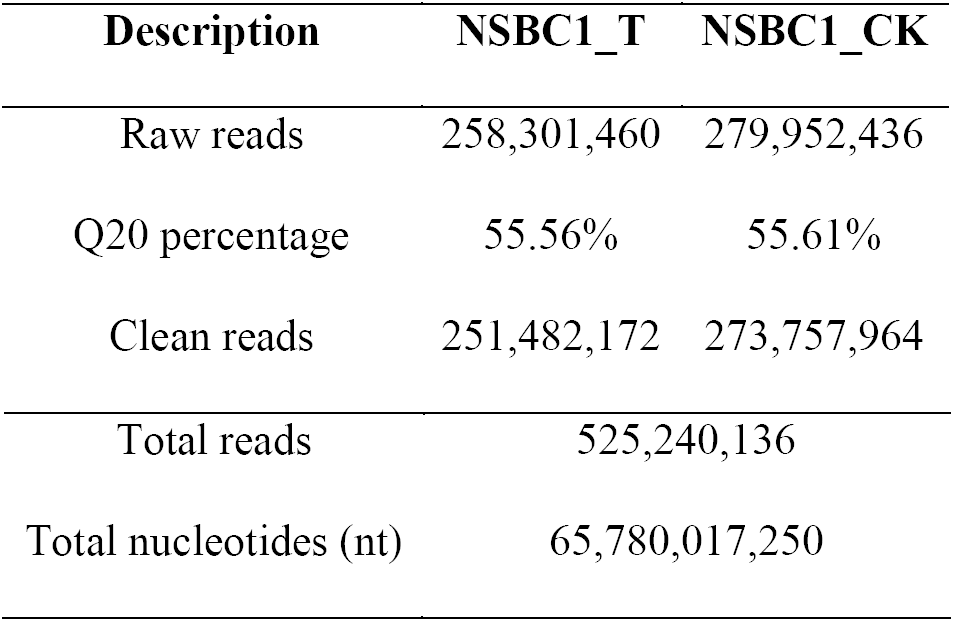
Overview of the transcriptome sequencing and *de novo* assembly results.

| Description | NSBC1_T | NSBC1_CK |
| --- | --- | --- |
| Raw reads | 258,301,460 | 279,952,436 |
| Q20 percentage | 55.56% | 55.61% |
| Clean reads | 251,482,172 | 273,757,964 |

|  |  |
| --- | --- |
| Total reads | 525,240,136 |
| Total nucleotides (nt) | 65,780,017,250 |

**Table 2.** Genome mapping summary of all samples

| Sample | Total Clean Reads | Total Mapping Ratio |
| --- | --- | --- |
| NSBC1_CK1-1 | 49273558 | 76.43% |
| NSBC1_CK1-2 | 51677702 | 77.54% |
| NSBC1_CK2-1 | 47765358 | 71.71% |
| NSBC1_CK2-2 | 44108084 | 75.97% |
| NSBC1_CK3-1 | 42718254 | 77.39% |
| NSBC1_CK3-2 | 44409480 | 76.30% |
| NSBC1_T1-1 | 44375910 | 77.42% |
| NSBC1_T1-2 | 42446806 | 78.30% |
| NSBC1_T2-1 | 23302262 | 78.14% |
| NSBC1_T2-2 | 23302264 | 77.40% |
| NSBC1_T3-1 | 42487034 | 76.63% |
| NSBC1_T3-2 | 41604074 | 77.65% |

**Fig 1.**
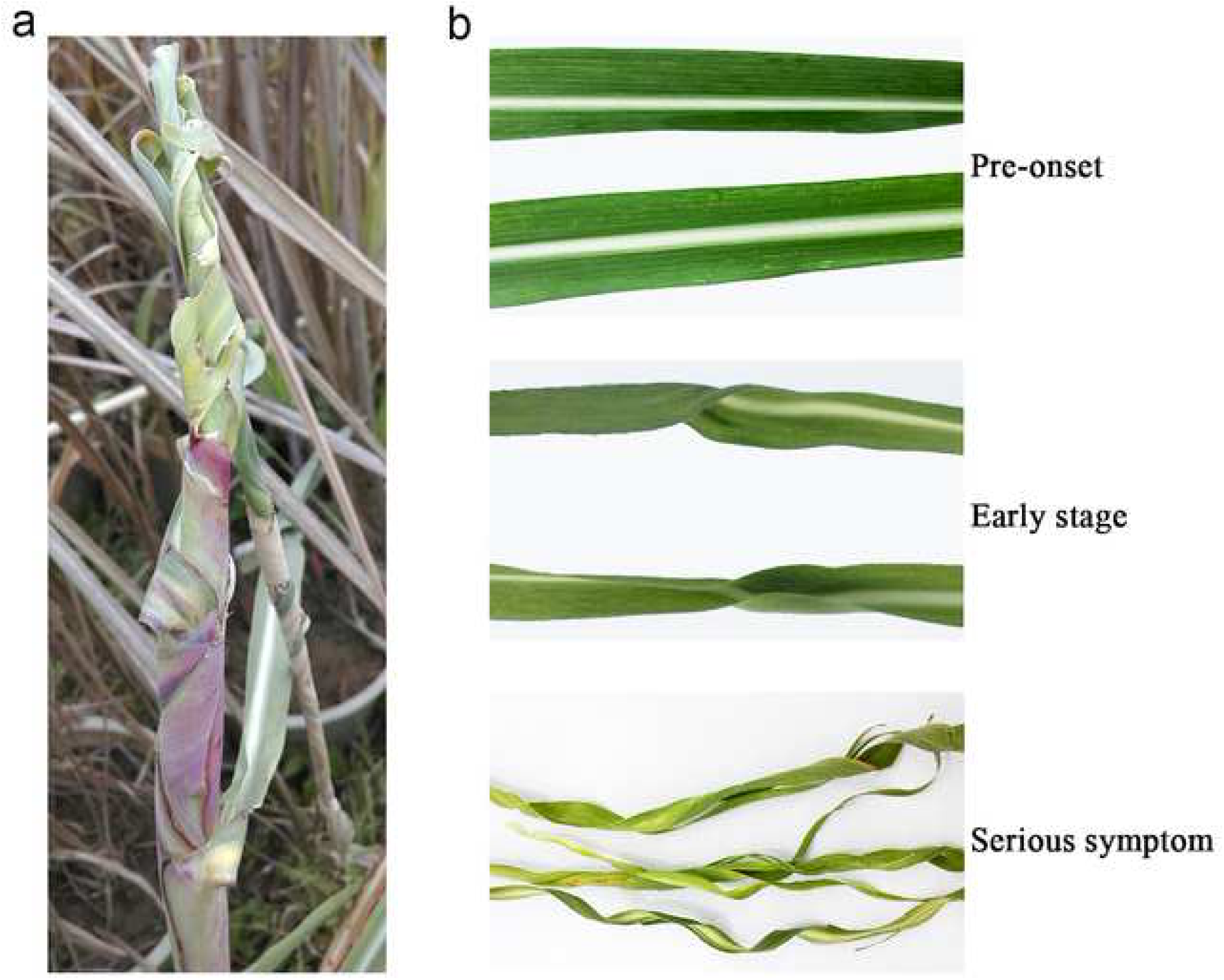
The susceptible sugarcane clone H3-8 under twisted leaf disease stress. (a) Twisted leaf disease of H3-8 was observed in the field. (b) Phenotype of three stages of Phoma sp. infection. From top to bottom: pre-onset, early, and serious symptom stages.

### Gene expression statistics

To elucidate gene expression profile of un-susceptible and susceptible cultivars, we annotated the aligned reads according to the reference genome. The result showed that there were 91,386 genes expressed that included 65,391 known genes and 25,995 novel genes. In the 96,101 annotated transcripts, 53,481 transcripts with novel alternative splicing subtypes encode known proteins, 27,151 transcripts were defined as novel protein coding genes, and 15,469 transcripts were classified into long non-coding RNAs. Details of each sample were exhibited in Table S1 and Figure S1.

Based on the Pearson correlation coefficient of genes expression of each sample, we obtained the correlation heatmap of all samples (Fig. 2a). The samples from the same cultivar conditioned high correlation Pearson value. Of course, two biological repeats of each stage clustered together in the Cluster Dendrogram (Fig. 2b). That means two biological repeats of each stage were consistent and satisfied for further analysis.

**Fig 2.**
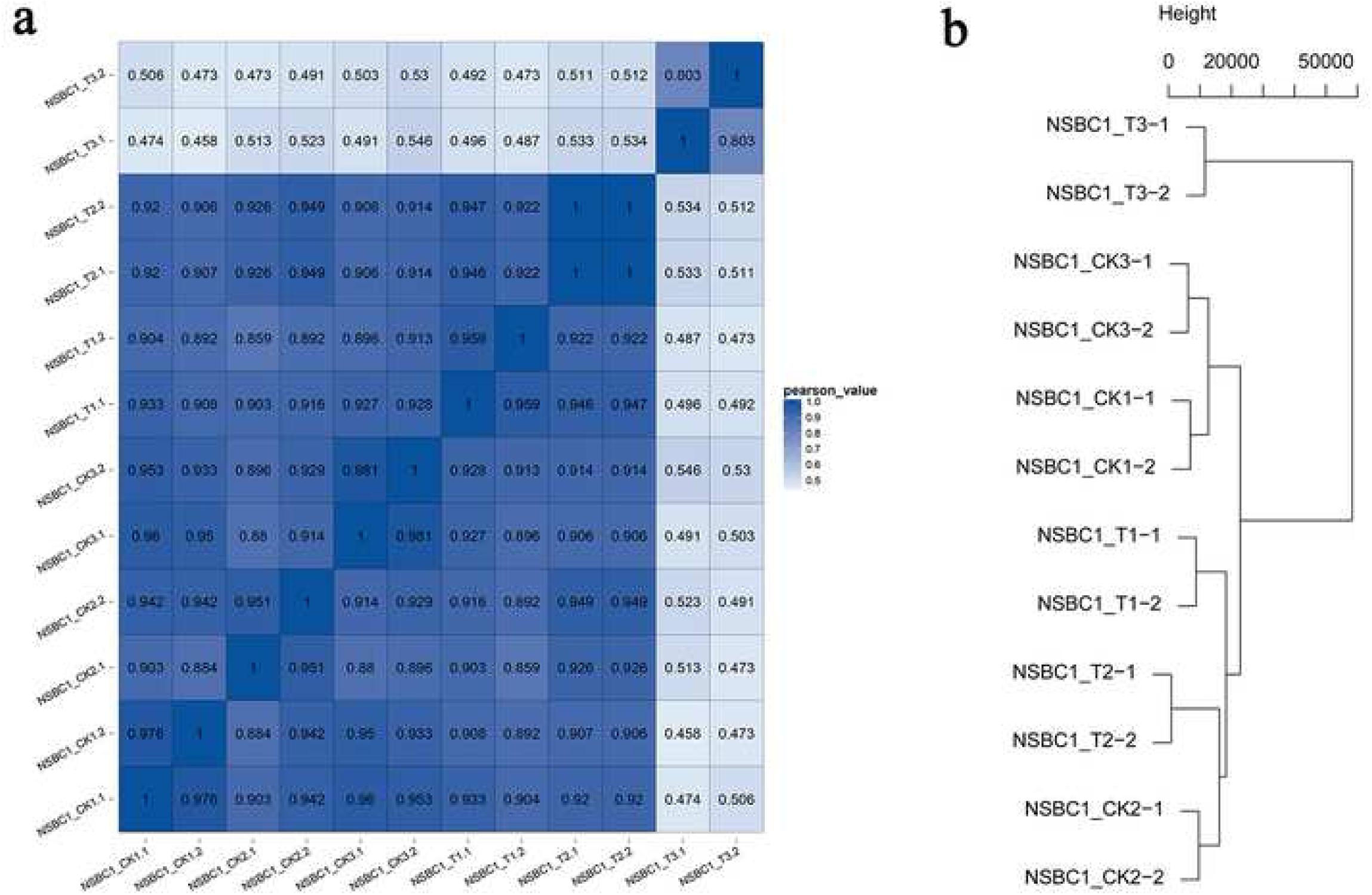
Correlation analysis of all samples. (a) Heatmap was constructed by the Pearson correlation coefficient of genes expression. (b) Cluster analysis of all samples based on genes expression quantity.

### Differentially Expressed Genes (DEGs) in Un-susceptible and Susceptible Cultivars

The significance of the gene expression difference was determined according to the threshold of FDR (False Discovery Rate) < 0.05 and |log_2_ FC (Fold Change) | >1. In the comparison to the un-susceptible clone H3-19, the genes expression of susceptible clone H3-8 showed 8,408 up-regulated and 7,834 down-regulated in pre-onset stage, 7,334 up-regulated and 5,558 down-regulated in early stage, and 24,456 up-regulated and 10,468 down-regulated in serious symptom stage. (Fig. 3). The Gene Ontology (GO) analysis classified DEGs into three main categories: biological function, cellular component, and molecular function (Fig. 4). DEGs mainly accumulated in the cellular process, metabolic process, and biological regulation in the biological function. The cell, cell part, and organelle conditioned most enriched DEGs in the cellular component. In the term of molecular function, DEGs were mainly distributed in catalytic activity and binding. Most enriched GO term are corresponding to various aspects of activated metabolism (e.g. “metabolic process” and “catalytic activity”) and stress response (e.g. “biological regulation” and “organelle”). We also performed a BLAST (The Basic Local Alignment Search Tool) (E-value <0.00001) analysis of the DEGs against the KEGG (Kyoto encyclopedia of genes and genomes) database. The DEGs were mainly enriched in metabolic pathways, biosynthesis of secondary metabolites, and plant-pathogen interaction (Fig. S2). These are consistent with the sugarcane activities in response to *Phoma* sp. infection. According to these findings, we analyzed specific pathways and genes related to disease defense.

**Fig 3.**
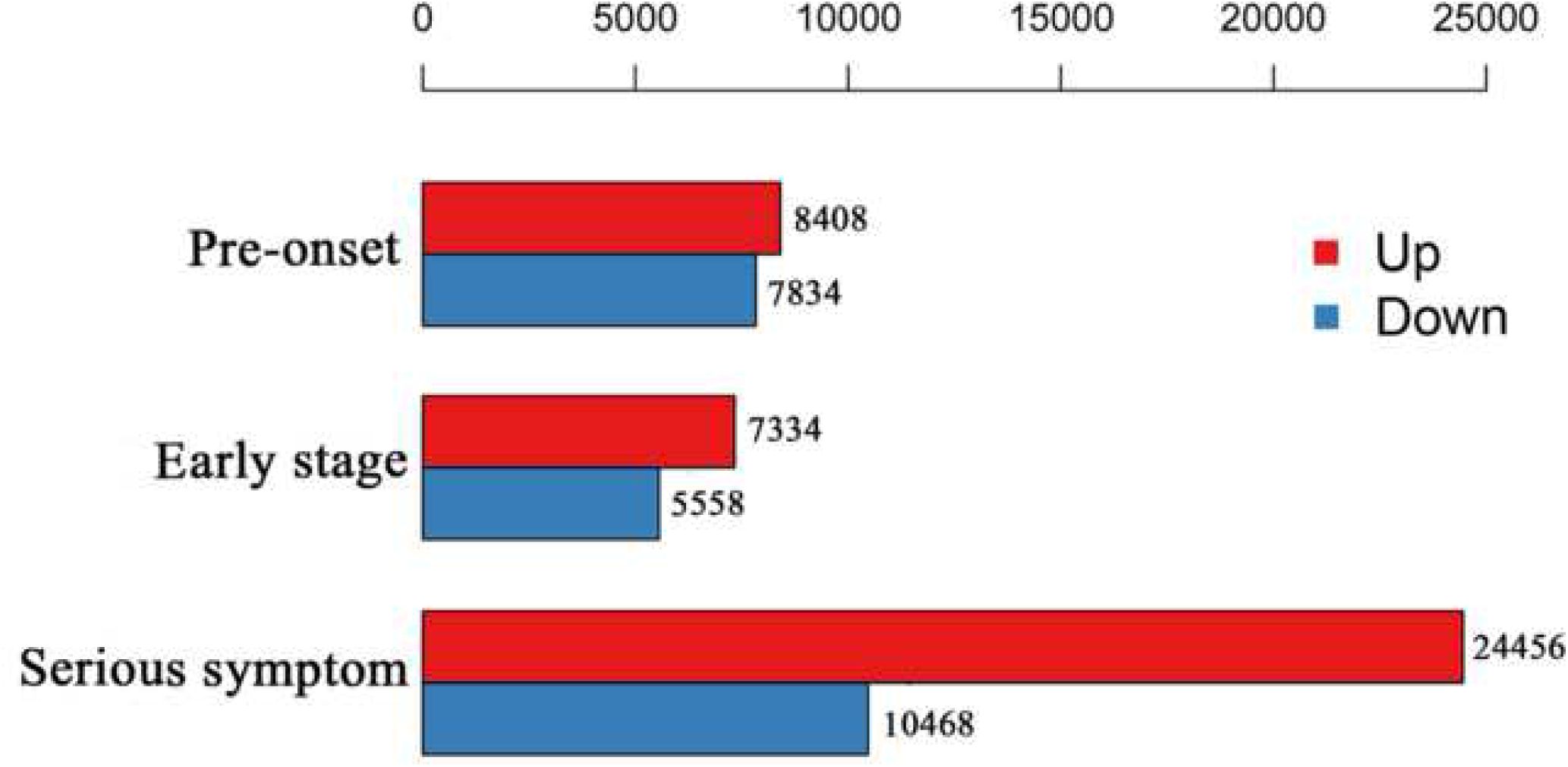
DEG analysis in the susceptible cultivars. Numbers of up- and down-regulated genes identified in the pre-onset, early, and serious symptom stages in susceptible genotypes.

**Fig 4.**
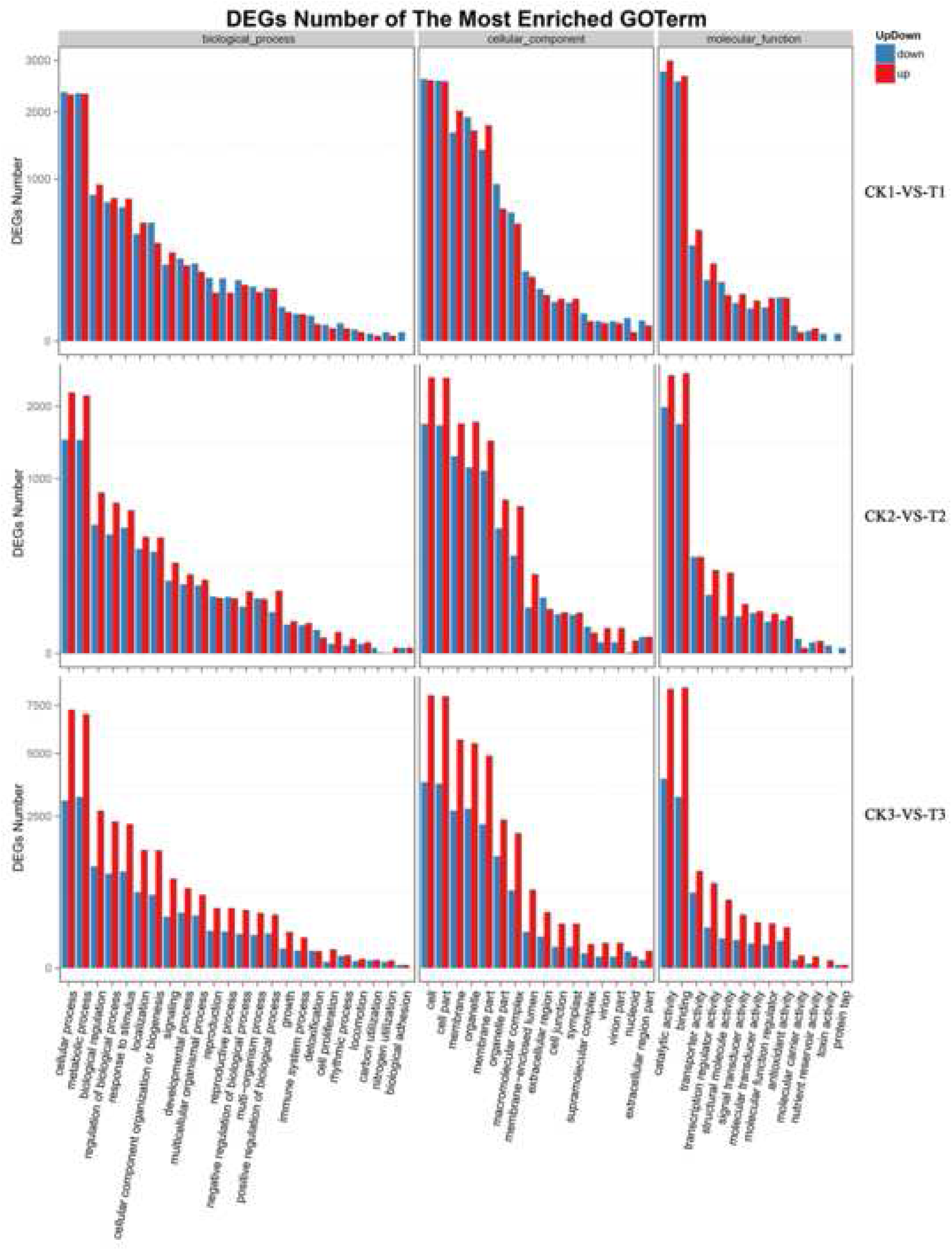
Gene ontology (GO) annotation for DEGs in three stages.

### Ingenuity Pathways Analysis

Plant hormones are the first reactors under environmental stress (Verma et al. 2016). Among the major hormones, abscisic acid (ABA), ethylene (ET), jasmonates (JA), and salicylic acid (SA) play essential roles in regulating plant response to biotic and abiotic stress (Bari and Ravindran 2009). The DEGs between the susceptible and un-susceptible cultivars provided us an opportunity to study how the hormone pathways response to twisted leaf disease. We analyzed the expression patterns of genes involved in hormone biogenesis in three different stage of *Phoma* sp. infection (Fig. 5; Table S1). As the main response hormones in biotic stress, the biosynthesis of SA, JA, and ET up-regulated in the susceptible cultivar (genes in green, brown, and blue lines). The SA, which is generally involved in the defense against biotrophic and hemi-biotrophic pathogens (Loake and Grant 2007), was enriched with high expression (∼4 folds up-regulated) of LOC_Os02g41630-D10and Sspon.04G0008070-3C (Fig. 5, green lines). JA and ET, which involve in the defense of necrotrophic pathogens and herbivorous insects (Gamalero and Glick 2012; Wasternack and Hause 2013), accumulated through up-regulating biosynthesis genes (Fig. 5, brown lines). ABA, auxin (IAA), and cytokinins (CK) biosynthesis genes also exhibited differential expressions in the susceptible cultivar, which resulted from hormone pathways crosstalk and secondary plant reflections (Robet-Seilaniantz et al. 2011). These results show that twisted leaf disease triggered the reaction of the hormones network.

**Fig 5.**
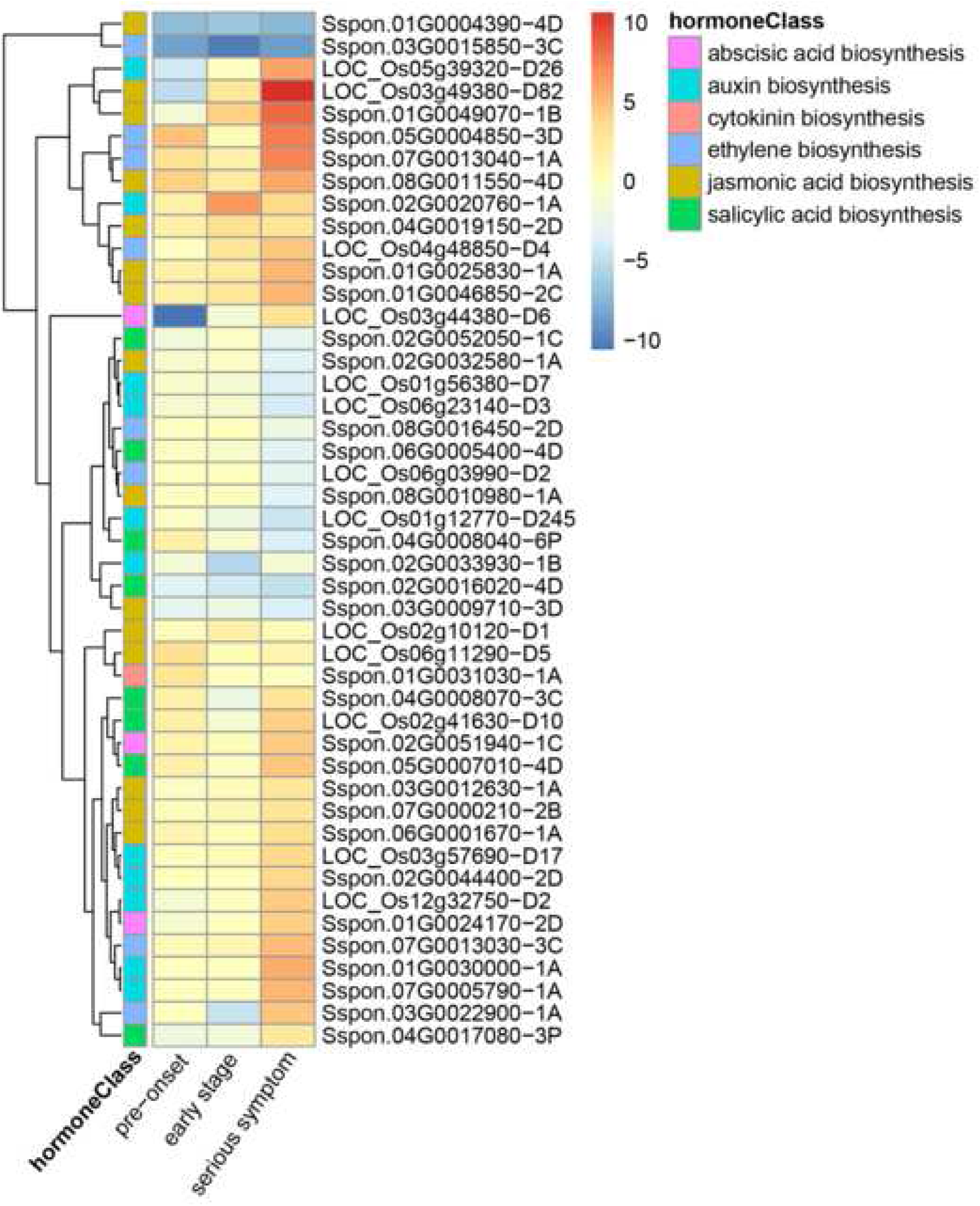
Ingenuity analysis of DEGs in hormone pathways during fungi infection process. Fold change (log_2_) in hormone biosynthesis genes at different stages of the infection process relative to the resistance condition.

Also, we analyzed the specific biosynthesis genes in SA, JA and ET pathway, respectively. In details, the phenylalanine ammonia-lyase, an enzyme of phenylalanine ammonium lyase (PAL) pathway in SA synthesis (Yokoo et al., 2018), was increased in expression at the pre-onset of *Phoma* sp. infection. The isochorismate synthase 1, which involved in the synthesis of SA (Yokoo et al., 2018), showed lower expression level in three stages of the susceptible cultivar (Table S1). In the JA pathway, 12-oxophytodienoate reductase, which catalyses the reduction of 10, 11-double bonds of 12-oxophytodienoate to 3-oxo-2-(2’-pentenyl)-cyclopentane-1-octanoic acid (OPC-8:0) (Tani et al., 2008), conditioned relative higher expression in the susceptible cultivar (Table S1). The aminotransferase, a gene faimly including VAS1 which participates in auxin and ethylene biosynthesis (Zheng et al. 2013), varied in expression at different stages of the twisted leaf disease. These findings supported that the infection of *Phoma* sp. induced the response of the hormones network.

### Evaluation of Differentially Expressed Disease-Resistant Genes

To defend against infection of pathogen, the expression of resistance (*R*) genes that encode proteins containing conserved nucleotide binding site plus leucine-rich repeat (NBS-LRR domain) are induced (Bakker et al. 2006). To elucidate the *R* genes involved in the twisted leaf disease, we annotated *R* genes in DEGs based on PRG (Pathogen receptor genes) database. (Table 3). There were 46%, 42%, and 26% *R* genes conditioned lower expression in pre-onset, early stage, and serious symptom of susceptible cultivar relative to the control, respectively. That means a part of *R* genes in the susceptible cultivar were null/retarded in the response of *Phoma* sp. infection.

**Table 3.** Classification of annotated *R* genes and representatives for ingenuity analysis.

| Class | Count | Representatives | Description |
| --- | --- | --- | --- |
| NL | 543 | Sspon.02G0020920-2B | Disease resistance protein (NBS-LRR class) family |
| RLP | 683 | Sspon.02G0041050-1B | Pyridoxal phosphate-dependent transferases superfamily protein |
| CNL | 242 | Sspon.05G0016050-1A | HOPZ-ACTIVATED RESISTANCE 1 |
| N | 309 | Sspon.08G0026530-1C | NB-ARC domain-containing disease resistance protein |
| TNL | 100 | Sspon.06G0034670-1D | TIR-NBS-TIR-TIR-WRKY type disease resistance protein |
| RLK | 7 | BGI_novel_G009740 | S-locus lectin protein kinase family protein |
| RLK-GNK2 | 94 | Sspon.01G0015790-1A | cysteine-rich RLK (RECEPTOR-like protein kinase) |
| CN | 67 | Sspon.04G0004450-1A | LRR and NB-ARC domains-containing disease resistance protein |
| Other | 40 | Sspon.01G0037110-1B | NADPH HC toxin reductase-like protein |
| T | 17 | Sspon.01G0011870-1A | NAC domain containing protein |
| Mlo-like | 21 | Sspon.01G0013680-3C | Seven transmembrane MLO family protein |
| L | 5 | Sspon.03G0025750-3D | Protein phosphatase 1 regulatory subunit |
| RPW8-NL | 4 | Sspon.02G0029950-1T | ADR1-like 1 disease resistance protein |

From the *R* genes expression profile, we selected out representative of each class to exhibit their expression patterns in three stages of *Phoma* sp. infection (Fig. 6). Sspon.06G0034670-1D, Sspon.04G0004450-1A and Sspon.02G0029950-1T dramatically decreased in three stages of *Phoma* sp. infection. Sspon.01G0011870-1A and Sspon.01G0013680-3C also showed lower expression level in the three stages. *R* genes like these genes were null function in the susceptible cultivar. Sspon.05G0016050-1A, Sspon.08G0026530-1C and Sspon.03G0025750-3D expressed lower at pre-onset and kept increasing in the later stages. That means *R* genes with such expression patterns in the susceptible cultivar were retarded in the defense of *Phoma* sp. infection. Some *R* genes like Sspon.02G0020920-2B, Sspon.02G0041050-1B, and Sspon.01G0037110-1B, conditioned higher expression or/and irregular expression patterns in three stages. These *R* genes could respond to *Phoma* sp. infection in the susceptible cultivar. Also, the sequence analysis of these representative genes, including alternative splicing and SNPs, didn’t offer clue for the structural mutation in the susceptible cultivar (data not shown). The expression level analysis of *R* genes revealed that the null and retarded response of *R* genes in the *Phoma* sp. infection, especially at the pre-onset stage might be one of the reasons that the H3-8 cultivar is susceptible to twisted leaf disease.

**Fig 6.**
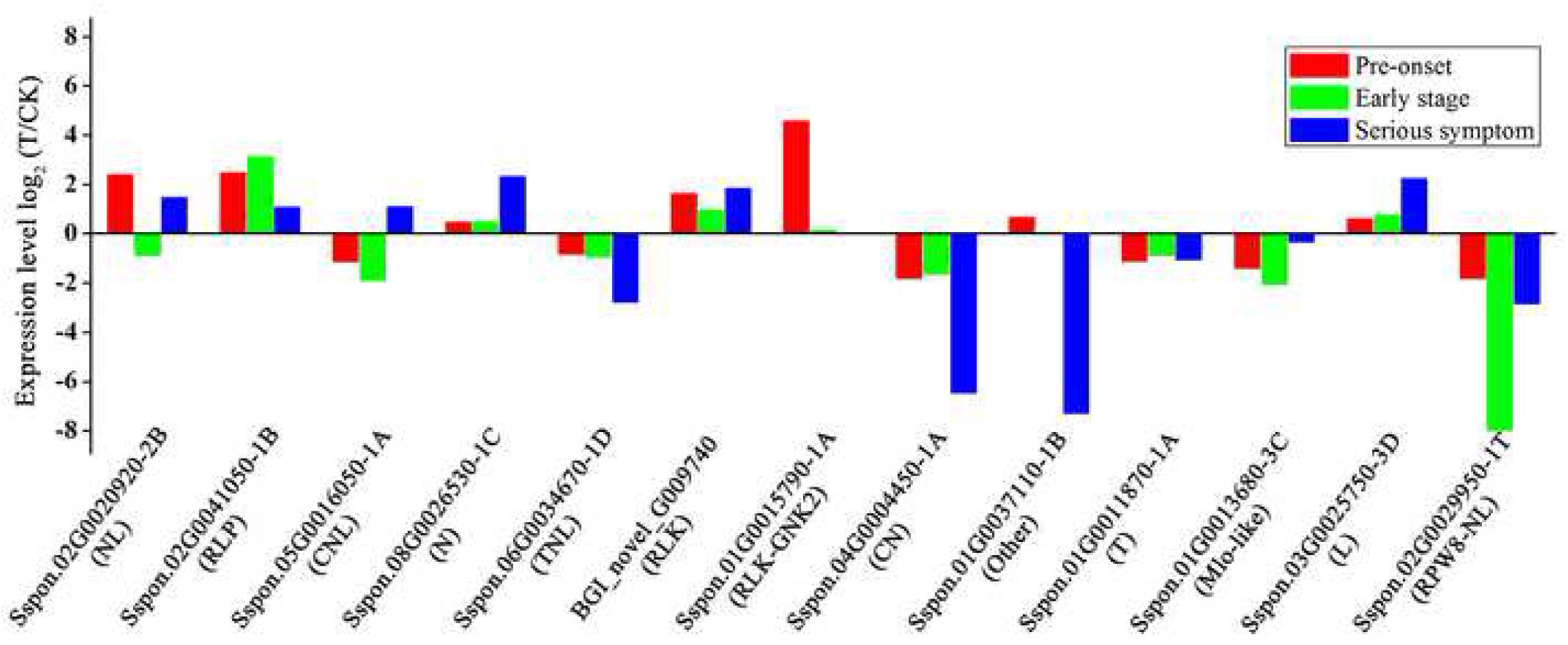
Differential expression analysis of specific *R* genes in the susceptible cultivar. Genes were selected out as representative based the tendency of each *R* gene class. Fold change (log_2_) was achieved from the FPKM value of T and CK.

## Discussion

The transcriptome, which is influenced by external environmental conditions, can effectively reveal the response mechanism of biotic and abiotic stress in plants (Wei et al. 2011). In our study, the natural infected cultivar H3-8 was defined as susceptible cultivar and the un-susceptible cultivar (H3-19) was used as control. With two biological repeats, we used transcriptome profiling to detect the genes associated with twisted leaf disease and obtained 65,780,017,250 bp data from three stages of the twisted leaf disease. Using the genome of *Saccharum spontaneum* L. as reference (Zhang et al. 2018), the average mapping ratio of twelve samples was 76.74%. The annotation of aligned reads showed 91,386 genes expressed (65,391 known and 25,995 novel), and 96,101 annotated transcripts, including 53,481 novel alternative splicing subtypes encode known proteins, 27,151 novel protein coding genes, and 15,469 long non-coding RNAs. With these data, we analyzed the DEGs, including hormone pathways and R genes, between the susceptible cultivar and the control. Also, these data provided a foundation for the further analysis of the twisted leaf disease, such as specific gene characterization, resistant genes selection, and immune network description.

The comparison between susceptible cultivar and the control identified 16,242, 12,892 and 34,924 DEGs in pre-onset, early stage and serious symptom stage, respectively (Fig. 3). These DEGs are involved in critical biological activities that are essential for disease resistance (Fig. 4). Ingenuity pathway analysis revealed hormone biosynthesis genes variations in the susceptible cultivar (Fig. 5). The SA and ET pathways were induced in the infection of *Phoma* sp. at the pre-onset stage. Along with the spread of the disease, JA and ET synthesis increased and maintained high levels in the serious symptom stages. As three major hormones involved in stress resistance, SA is a biotrophic and hemi-biotrophic pathogen-triggered signaling pathway, and JA combined with ET signaling pathways are induced by necrotrophic pathogens (Glazebrook 2005). Therefore, twisted leaf disease, caused by fungal pathogens, first triggers the SA pathway as response to pathogens infection. The high level of JA and ET in serious symptoms might partially result from tissue damage in disease. Other hormones, including ABA, auxin, and CK, crosstalk with major disease response hormones (SA, JA, and ET) and construct a network to respond to disease stress (Verma et al. 2016).

Plant disease resistance (*R*) genes are prevalent in all plant species and harbor a conserved LRR domain (Meyers et al. 1998). *R* genes are defined as gene-for-gene interaction and specifically recognize an avirulence protein encoded by a pathogen with a hypersensitive response (Flor 1956). In the *R* gene analysis of the susceptible cultivar, around 40% *R* genes, including those that play essential functions, were down-regulated, especially in pre-onset stage. For example, belongs to LRR and NB-ARC domains-containing proteins of which members participate in disease resistance (Fischer et al. 2016). is predicted to encode RGA protein which contains NB-ARC domain and functions disease resistance (Cesari et al. 2013). is descripted as *CRKs* (cysteine-rich protein kinase), which involves in the ROS signaling pathway and plays important roles in the elimination of fungal growth damage (Niina 2015). The low expression of these *R* genes in the twisted leaf disease would drag down the rapid response of *R* genes defensive system. Besides ∼40% down-regulated *R* genes, there were still a number of *R* genes conditioned normal expression level or even higher expression level in the susceptible cultivar. Unfortunately, these high expressed *R* genes were unable to eliminate the pathogens successfully. These increased expressions might be a result of universal reactions of sugarcane under pathogens stress rather than the specific response of *Phoma* sp. infection. Such phenomenon was commonly reported in the studies of other species (Heath 2000; van Loon 2015).

## Materials and Methods

### Plant Materials and Sampling

*Narenga porphyrocom*a, which is an important relative genus of sugarcane, was collected from a barren mountainous area in Guangxi province, China. We obtained the BC_1_ generation offspring of the *Narenga porphyrocom*a via crossing and backcrossing with sugarcane varieties several years. Among the BC_1_ offspring, H3-8 was susceptible to twisted leaf disease at the sugarcane elongating stage, whereas H3-19 was not. H3-8 was proven to affect stable twisted leaf disease occurrence after three years based on field and greenhouse observations.

Plantlets at the same growth stage were selected and planted in pots filled with a mixture of peat soil and washed sand, and grown in a greenhouse at the Sugarcane Research Institute, Guangxi Academy of Agricultural Sciences. The natural infected plants were selected and defined as the susceptible cultivator. The leaf samples of the susceptible sugarcane clone H3-8 were collected at three stages corresponding to the pre-onset (named NSBC_1__T1), early (named NSBC_1__T2), and serious symptom (named NSBC_1__T3) stages of the disease, respectively. The leaf samples of the un-susceptible sugarcane clone H3-19 were simultaneously collected, labeled NSBC_1__CK1, NSBC_1__CK2, and NSBC_1__CK3, respectively. Two biological repeats of each sample were collected. All samples were immediately frozen in liquid nitrogen and stored at –80°C.

### RNA Extraction and Quality Determination

The total RNA was extracted using TRIzol reagent (Invitrogen, Carlsbad, CA, USA) and treated with RNase-free DNase I and RNA integrity number (RIN > 8.0). RNA quality and quantity were verified using a NanoDrop 1000 spectrophotometer (Wilmington, DE, CA, USA) and an Agilent 2100 Bioanalyzer (Santa Clara, CA, USA) prior to the library construction. No sign of degradation was found.

### cDNA Library Construction and Sequencing

The poly(A) RNA was fragmented into approximately 300-nt fragments using RNA Fragmentation Reagents (Ambion, Austin, TX, USA). Using these short fragments as templates, the double-stranded cDNA was synthesized with random primers (Invitrogen, Carlsbad, CA, USA). The end repair of these cDNA fragments was subsequently performed with Klenow polymerase, T4 DNA polymerase, and T4 polynucleotide kinase (NEB, Ipswich, MA, USA). Illumina adapters (containing primer sites for sequencing and flow cell surface annealing) were ligated to the end-repaired fragments using T4 DNA ligase (Invitrogen, Carlsbad, CA, USA). The products were enriched for the cDNA fragments using a Qiaquick Gel Extraction Kit (Qiagen, Duesseldorf, Germany) and amplified with polymerase chain reaction (PCR) to prepare the sequencing library. Agilent 2100 Bioanalyzer (Agilent, Beijing, China) was used to detect the quantity and quality of the cDNA. Then, the paired-end RNA-seq libraries were prepared following Illumina’s protocols and sequenced on the Illumina HiSeq 2500 platform (Illumina, San Diego, CA, USA). Sequencing was performed in a flowcell on an Illumina HiSeq 2500 sequencer using the TruSeq Paired-End Cluster Kit v3 (Illumina PE-401-3001) and the TruSeq SBS HS Kit v3200 cycles (Illumina FC-401-3001), generating 2 × 125 bp.

### Data Filtering and Genome Mapping

The raw reads were filtered according to the following criteria: (1) reads containing adaptor, (2) reads with unknown nucleotides larger than 5%, and (3) low-quality reads (the rate of reads with a quality value of ≤10 was more than 20%. The clean data was mapped to the genome of *Saccharum spontaneum* L. (Zhang et al. 2018) via HISAT (v2.0.4; --phred64 --sensitive --no-discordant --no-mixed -I 1 -X 1000) (Kim et al. 2015). The transcripts were re-constructed via StringTie and predicted using Cuffcompare tool in Cufflink (Trapnell et al., 2010; Pertea et al., 2016).

### Differential Gene Expression

The DEGs analysis was performed as the description of DEGseq (Wang et al. 2010) and corrected the *P*-values to *Q*-values based on Benjamini-Hochberg method (Benjamini and Hochberg 1995) and Storey-Tibshirani method (Storey and Tibshirani 2003). GO (gene ontology) terms were assigned based on the best-hits BLASTx resulted from NR alignments that were derived from Blast2GO (v2.5.0) against GO database (release-20120801). The DEGs were aligned against KEGG (Kyoto Encyclopedia of Genes and Genomes) by BLASTx package with threshold of E-value <=10-5.

### Ingenuity Analysis of DEGs

The rice hormones proteins were used as reference to identify the hormone genes in sugarcane (Cohen et al. 2017). The longest transcript of genes was selected as representative for analysis. All predicted protein sequences were aligned against the sugarcane gene models by BLASTp (Identity> 50%; Coverage>50%; E-value: 1×10^7^), and defined as homologs of hormones biosynthesis genes. Also, we aligned all predicted hormone genes against the sugarcane genome via tBLASTn (E-value: 1E-10), and used GeneWise (v2.2.0) to identify the structure of candidate hormone genes. Among these genes, the ones with non-significant different expression in three stages were eliminated. The final candidates were aligned against KEGG database (version:84). The log fold of each gene was obtained from the case-control group. The patterns of the gene in each group were compared. Similarly, predicted proteins were aligned against the PRG database using BLASTp. The results were filtered (E-value: 1e10), and the best hit of predicted protein was retained. The fpkm (Reads per kilobase of exon per million reads mapped) of each predicted resistant gene in the case sample was filtered out and a downstream analysis was performed.

## Abbreviations

NR: non-redundant protein sequences in NCBI
COG/KOG: clusters of orthologous group
KEGG: Kyoto encyclopedia of genes and genomes databases
TGICL: TIGR gene indices clustering tools
Blast: Basic Local Alignment Search Tool

## Acknowledgments

We thank Mick Rose in Primary Industry Department of Australia for the assistance of writing paper.

## Supplementary Materials

**Supplemental Figure 1:**
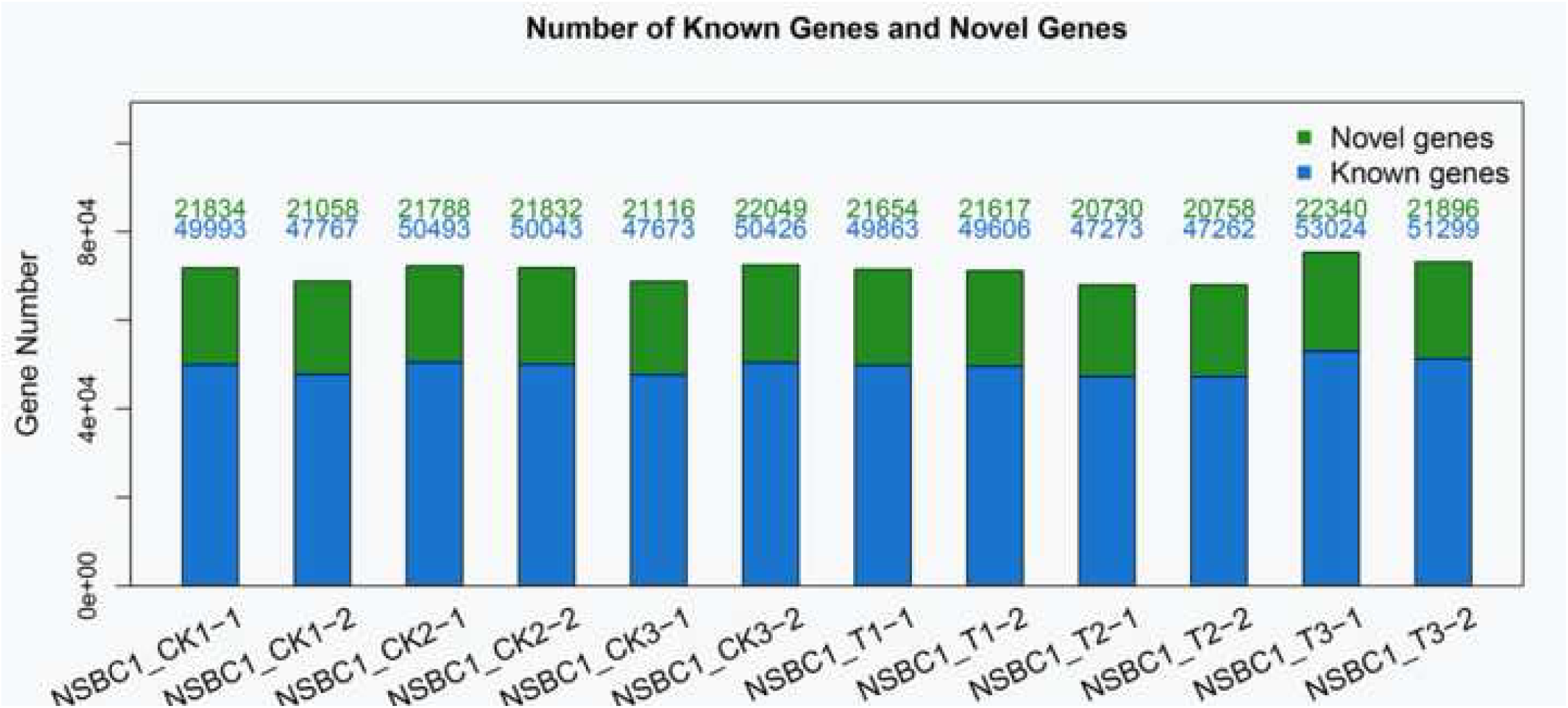
Details of gene annotation of all samples.

**Supplemental Figure 2:**
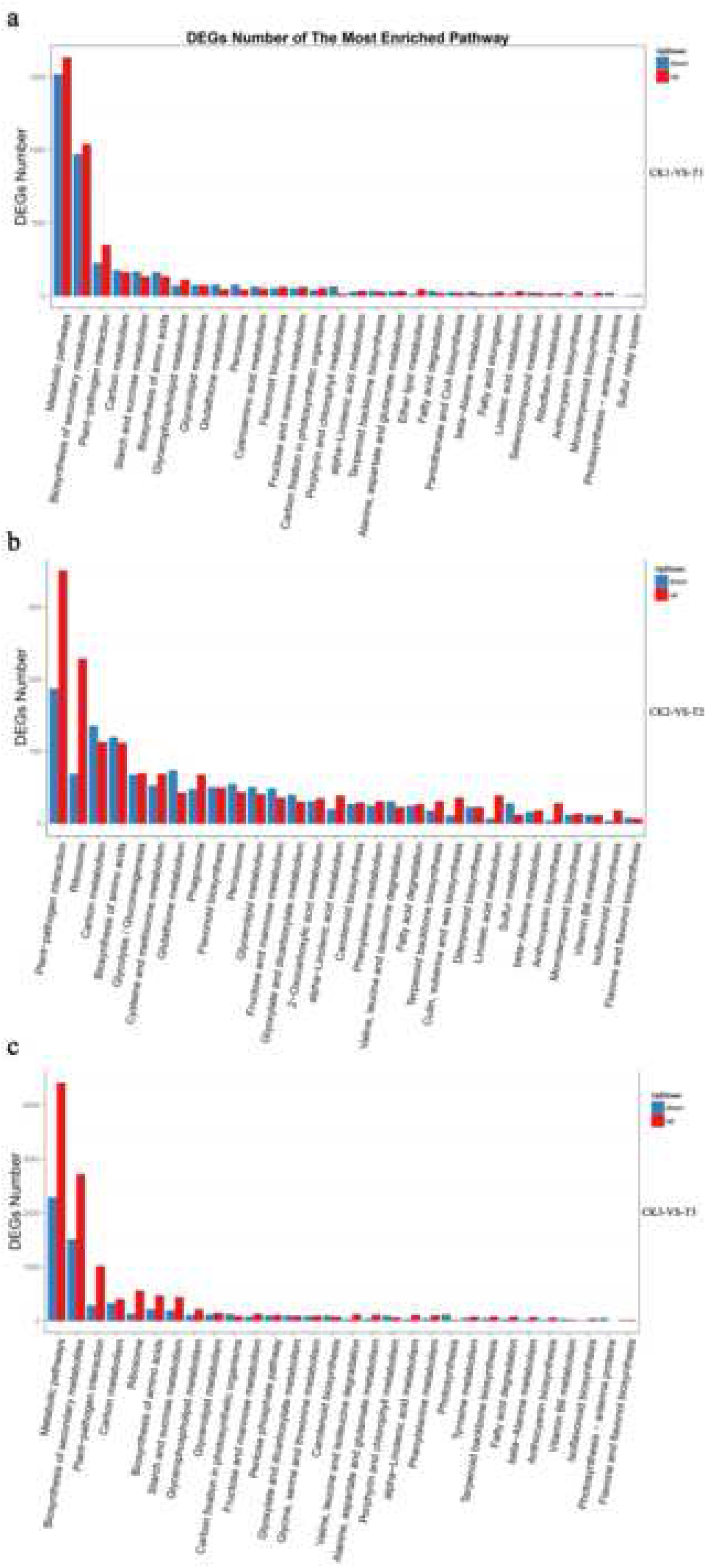
Annotation of DEGs against KEGG database.

Supplemental Table 1: Details of DEGs of hormones pathways analysis in susceptible relative to the control

## Author Contributions

Hongwei Tan, and Xihui Liu conceived and designed the experiments. Jinju Wei and Huiping Ou performed the experiments. Xiaoqiu Zhang, Ronghua Zhang, Hui Zhou, Yiyun Gui, Haibi Li, Yangrui Li, Rongzhong Yang and Dongliang Huang performed cross and backcross of *Narenga porphyrocoma*, planted sugarcane and collected samples. Zhihui Xiu, Junhui Chen and Huayan Jiang analyzed the data. Zhihui Xiu and Xihui Liu wrote the manuscript. All authors have read and approved the final manuscript.

## Conflicts of Interest

The authors declare no conflict of interest.

The data that support the findings of this study have been deposited in the CNSA (https://db.cngb.org/cnsa/) of CNGBdb with accession code CNP0000220, and NCBI with accession code SAMN05428728 to SAMN05428739.

